# Automated neuropil segmentation of fluorescent images for Drosophila brains

**DOI:** 10.1101/2024.02.03.578770

**Authors:** Kai-Yi Hsu, Chi-Tin Shih, Nan-Yow Chen, Chung-Chuan Lo

## Abstract

The brain atlas, which provides information about the distribution of genes, proteins, neurons, or anatomical regions in the brain, plays a crucial role in contemporary neuroscience research. To analyze the spatial distribution of those substances based on images from different brain samples, we often need to warp and register individual brain images to a standard brain template. However, the process of warping and registration often leads to spatial errors, thereby severely reducing the accuracy of the analysis. To address this issue, we develop an automated method for segmenting neuropils in the *Drosophila* brain using fluorescence images from the *FlyCircuit* database. This technique allows future brain atlas studies to be conducted accurately at the individual level without warping and aligning to a standard brain template.

Our method, LYNSU (Locating by YOLO and Segmenting by U-Net), consists of two stages. In the first stage, we use the YOLOv7 model to quickly locate neuropils and rapidly extract small-scale 3D images as input for the second stage model. This stage achieves a 99.4% accuracy rate in neuropil localization. In the second stage, we employ the 3D U-Net model to segment neuropils. LYNSU can achieve high accuracy in segmentation using a small training set consisting of images from merely 16 brains. We demonstrate LYNSU on six distinct neuropils or structure, achieving a high segmentation accuracy, which was comparable to professional manual annotations with a 3D Intersection-over-Union(IoU) reaching up to 0.869.

Most notably, our method takes only about 7 seconds to segment a neuropil while achieving a similar level of performance as the human annotators. The results indicate the potential of the proposed method in high-throughput connectomics construction for *Drosophila* brain optical imaging.

## Introduction

Over the past few decades, neuroscience has evolved from a discipline primarily dependent on experimental biology into an interdisciplinary field of study^1,2^. Notably, breakthroughs in imaging applications, such as high-resolution neural imaging techniques, including fluorescence microscopy and electron microscopy^3–9^, have been significant. These advancements have enabled us to observe neural tissues and circuits with unprecedented speed and resolution. In this context, analyzing variations in brain structures across different individuals or the expression intensity of genes in various brain regions has become a crucial research direction^10–13^. Consequently, developing an automated and high-throughput method for this analysis has become particularly important^14–16^.

Indeed, a number of studies have developed algorithms of brain region segmentation for CT and MRI images^17–25^. However, a similar method for optical images of *Drosophila* brain has not yet been developed. The Drosophila brain contains 58 neuropils^26^, and each neuropil, if segmented manually, would take up to four hours on average. Therefore, segmenting every neuropil manually for each optical images of the *Drosophila* brain is impractical. Instead, a common practice is to warp and align optical brain images obtained from different individuals to a standard brain template^26,27^ which contains segmented brain regions and perform the subsequent analysis in the warped brain images. While this method facilitates subsequent statistical analysis and interpretation, it also introduces spatial errors^29–31^. These inaccuracies could impact the precise interpretation of neural circuits, thereby reducing the reliability of research findings. To address this issue, one should segment neuropils directly in the original images to avoid errors introduced during the alignment process.

Developing more accurate and efficient image analysis algorithms becomes exceptionally crucial. This involves improving the accuracy of segmentation and alignment and considering computational costs and time efficiency to meet the needs of large-scale, high-throughput studies. Simultaneously, segmenting Neuropils directly from the original data can avoid errors introduced during the alignment process, thus enhancing the accuracy and reliability of the analysis.

In the present paper, we introduce a novel computational method, LYNSU (Locating by YOLO and Segmenting by U-Net), specifically designed for segmenting neuropils in *Drosophila* brain fluorescence images. Our method is divided into two stages based on a detection-led segmentation workflow. In the first stage, we detect the location of the neuropil of interest using the YOLOv7^32^ model, renowned for its exceptional inference speed and rapid convergence during training. These characteristics make YOLOv7 particularly suitable for high-throughput task requred for large databases like *FlyCircuit*^26^, significantly reducing computational time for detecting neuropils. In the second stage, we segment the neuropil in the bounding box defined by YOLOv7 using the 3D U-Net^33–35^ model, a deep learning model specifically designed for three-dimensional image segmentation.

The innovation of this method lies in its combination of advanced object detection technology with specialized image segmentation algorithms, achieving efficient and accurate segmentation. Through LYNSU, we anticipate more in-depth and detailed analyses of *Drosophila* brain structures, bringing new insights to neuroscience. This study demonstrates the potential applications of computational methods in biomedical image analysis and provides a reliable reference model for future high-throughput image analyses.

## Results

The proposed method, LYNSU, is designed based on a two-stage approach with an aim to achieve high-precision and high-speed brain segmentation (Figure 1). The goal of the first stage is to locate the target neuropils quickly and accurately. We first convert a 3D brain image stack into a 2D image by calculating the maximum and average brightness along the Z-axis direction, resulting in two 2D images. This ensures that the YOLOv7 model is exposed to a variety of brightness information in the images, enhancing its ability to identify the target neuropil. In the training phase, the YOLOv7 model is trained on human-labeled bounding boxes encompassing the target neuropil. In the test phase, the YOLOv7 model can rapidly generate a Region of Interest (ROI) containing the target brain neuropil for each 2D image. The goal of the second stage is to segment the target neuropil from the 3D image stack within the ROI generated from the first stage. In the graining phase, the 3D U-Net model is trained on human-segmented 3D image stacks. These 3D stacks are sliced into multiple cubes through an overlapping sliding method for data augmentation. We chose the 3D U-Net model for this delicate segmentation task, as this model can accurately capture the complex three-dimensional structure of neuropils and further enhance segmentation precision. Finally, the workflow achieves extremely high operational efficiency and accuracy in the testing phase.

**Fig. 1.**
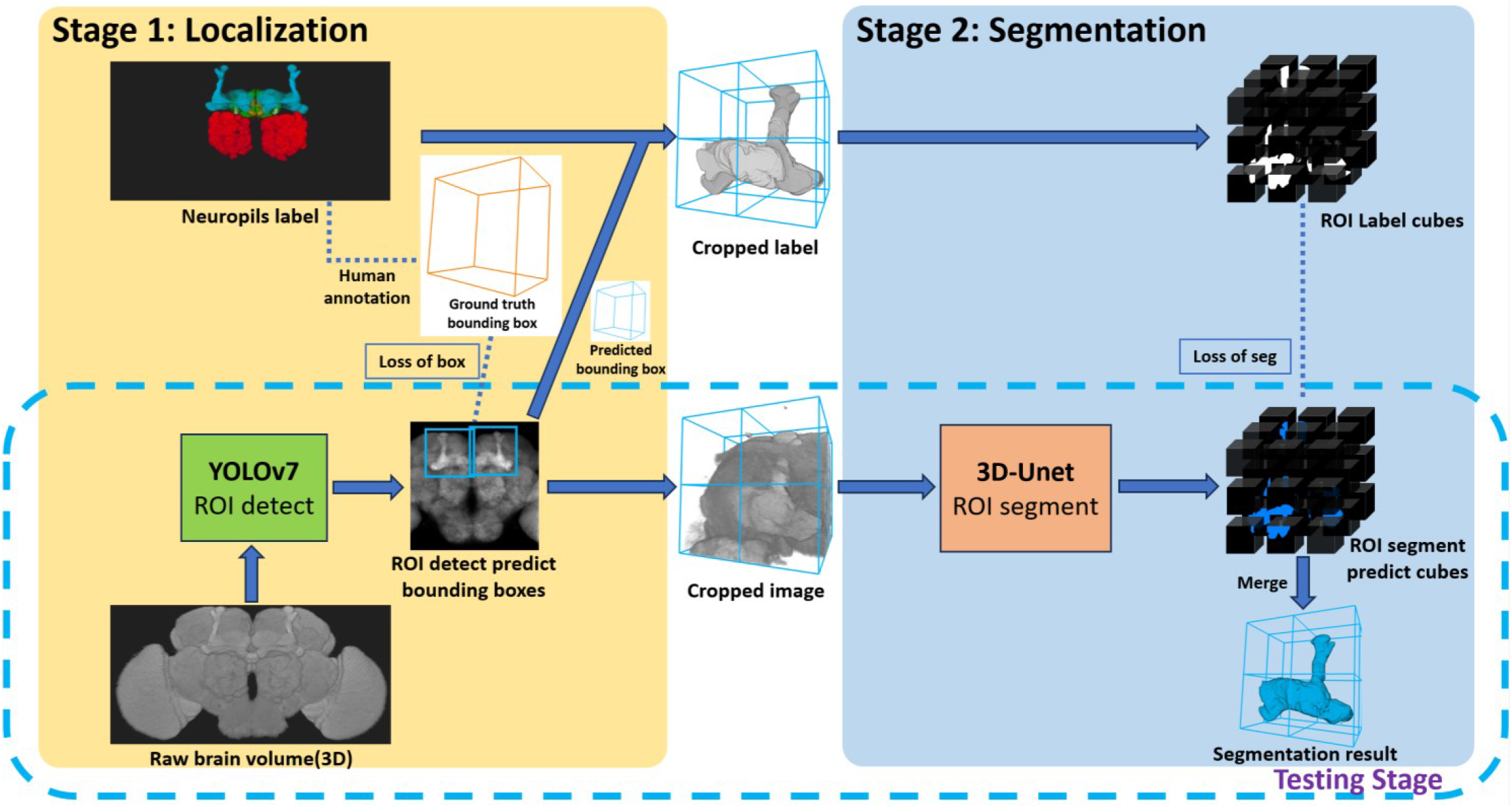
The LYNSU workflow. In Stage 1, rapid localization of neuropils is achieved using 2D brain projections and YOLOv7, resulting in the generation of ROIs encompassing complete neuropils. In Stage 2, 3D images are extracted from these ROIs and precisely segmented using the 3D U-Net model. The testing phase only required the workflow in the dashed box.

We first demonstrate the result of mushroom body (MB) segmentation with brain samples from three test sets: Gad1-F-400041, VGlut-F-800014, and Trh-F-200069. It is important to note that these three samples are 3D images not previously used by the model and come from different random splits of the dataset. We successfully achieved high-precision segmentation of through the model’s inference process (Figure 2). These two-dimensional sections, compared to the corresponding ground truth, clearly demonstrate that the MB in three different brain samples were precisely segmented by LYNSU. We further demonstrate the segmentation results of five other neuropils or brain structure, AL (Antennal Lobe), CAL (Calyx), FB (Fan-shaped Body), EB (Ellipsoid Body), and PB (Protocerebral Bridge), from different sample brains in the test set (Figure 3).

**Fig. 2.**
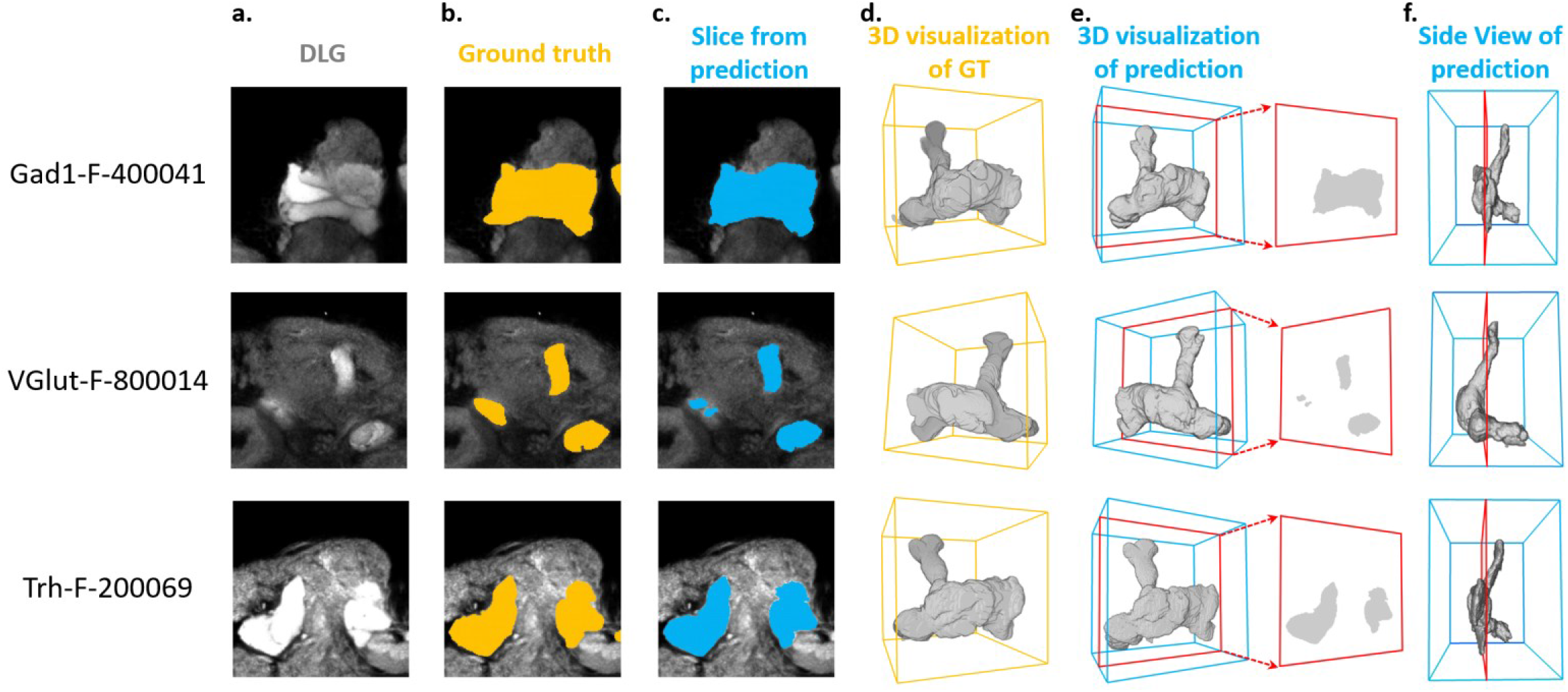
Segmentation of the MB (Mushroom Body) in three brain samples: Gad1-F-400041, VGlut-F-800014, and Trh-F-200069 (from top to down). **(a)** a slice of the anti-DLG image from the sample brain, **(b)** human segmentation as the ground truth, **(c)** segmentation by the proposed method (LYNSU), **(d)** 3D reconstruction of the ground truth segmentation, **(e)** 3D reconstruction of the LYNSU segmentation with the slice as shown in **(c)** and **(f)** the side view of the LYNSU segmentation. The red frames indicate the location of the slice. The display of different layer slices from these three MB samples showcases the precision of our segmentation method.

**Fig. 3.**
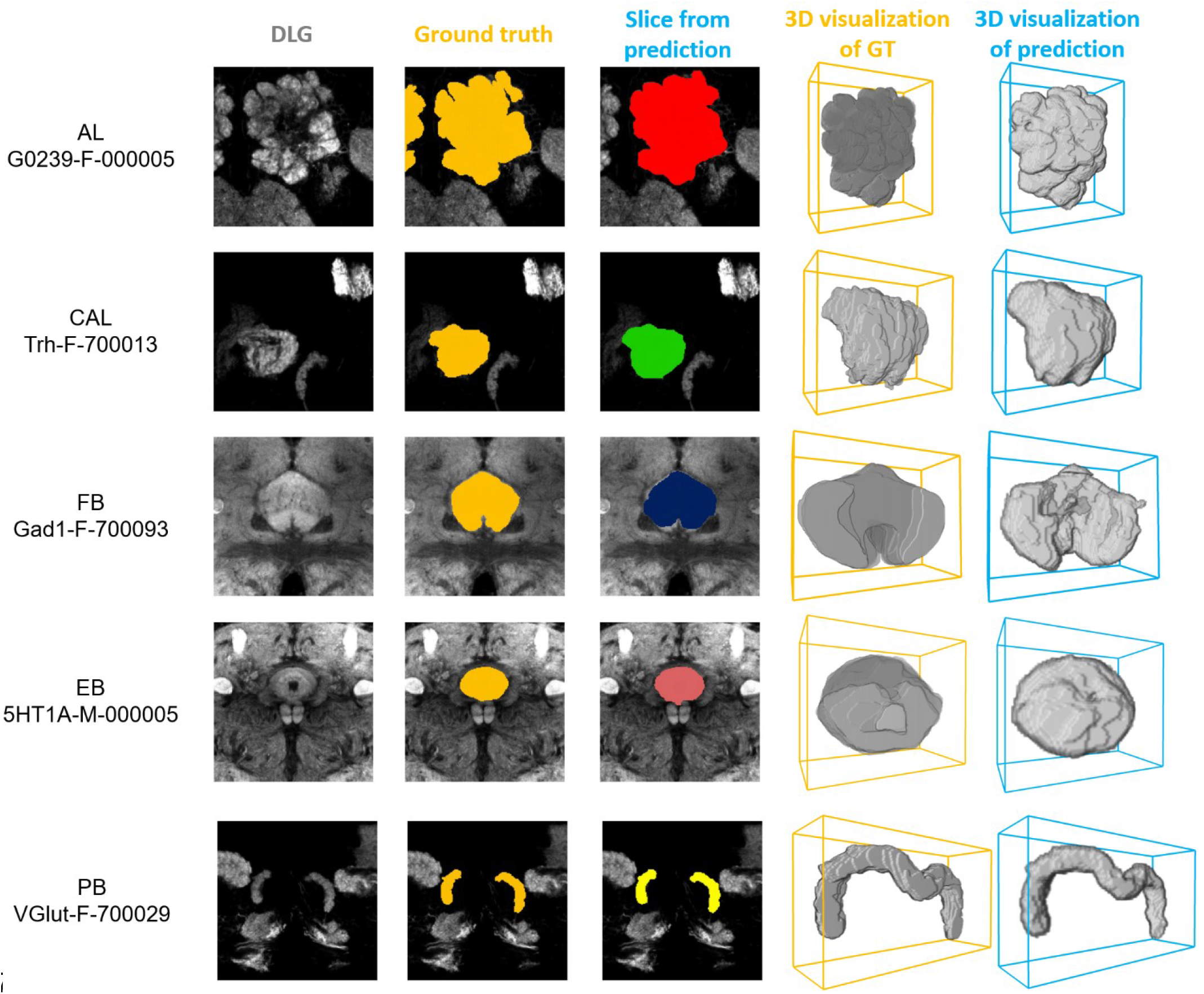
The segmentation results for five neuropils or brain structures: AL, CAL, FB, EB, and PB. From left to right: a slice of the anti-DLG image from the sample brain, the ground truth (GT), LNYSU segmentation, 3D reconstruction of GT and 3D reconstruction of LNYSU segmentation.

We systematically compare LYNSU to three other mainstream segmentation algorithms without using YOLO: FCN^36^, 2D U-Net, and 3D U-Net. First, visual inspection of the 2D slices indicates that LYNSU produce more precise neuropil boundaries than other algorithms (Figure 4a). Next, we conducted a quantitative evaluation and using four commonly used metrics: Recall, Precision, F1 score, and 3D-IoU. The LYNSU model outperformed the other three algorithms (Figure 4b). Specifically, LYNSU model performs much better than FCN and 2D U-Net by a large margin in all metrics, and is also much better than 3D U-Net in three out of four metrics.

**Fig. 4.**
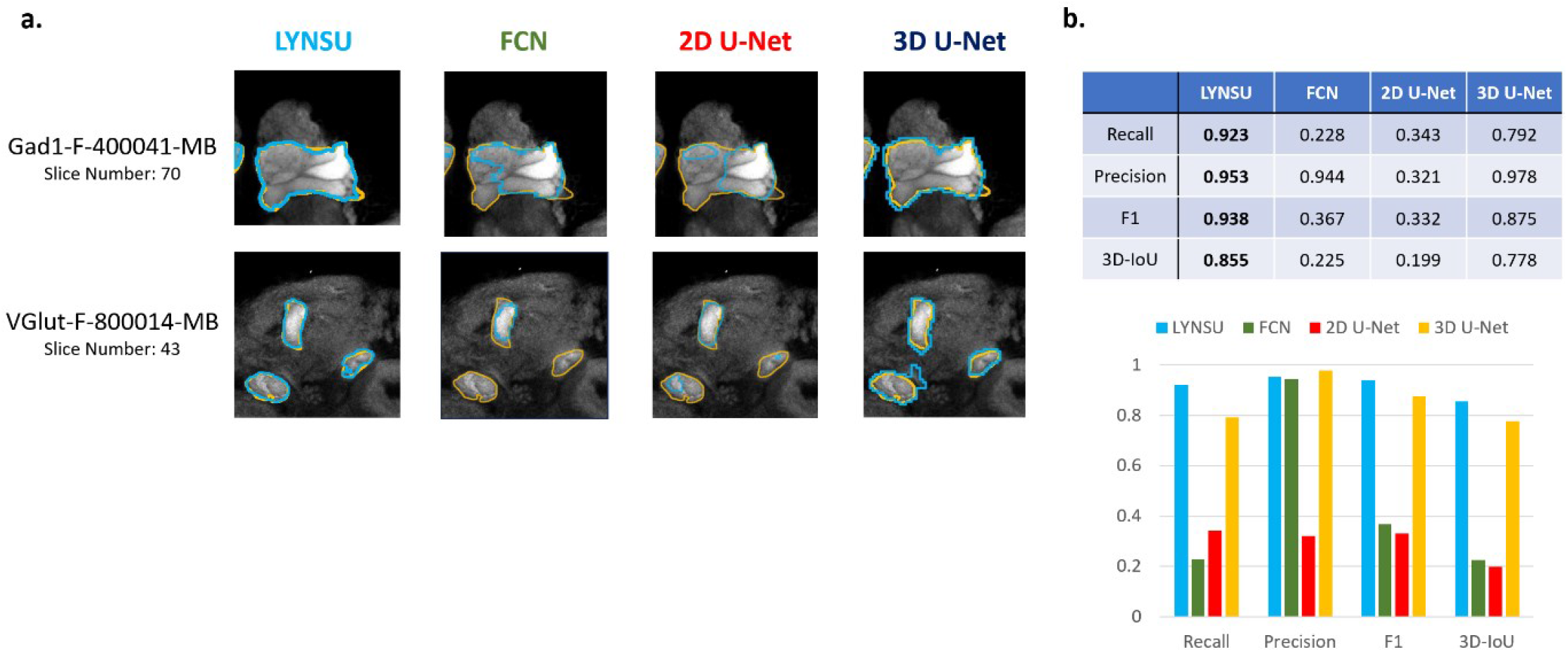
Performance comparison between LYNSU, FCN, 2D U-Net, and 3D U-Net based on the images from the *FlyCircuit* dataset. (a) Selected 2D slices from two sample brains (top and down) demonstrate superior boundary delineation by LYNSU. Orange: ground truth. Blue: predicted boundaries by each algorithm. (b) Quantitative metrics, including Recall, Precision, F1, and 3D-IoU, indicate much better performance of LYNSU than other algorithms.

We conducted ten random splits of the dataset with 16 brains assigned as the training set and two brains as the test set. We test the performance of LYNSU against 3D U-net (without YOLO) for all six neuropils or brain structures (AL, MB, CAL, FB, EB, PB), and LYNSU achieve higher 3D IOU score than 3D U-net for all neuropils or brain structures (Figure 5) with an average 3D IOU exceeding 0.833 comparing to 0.763 for 3D U-net. These results strongly validate the capability of LYNSU in segmenting various brain regions of the *Drosophila* brain.

**Fig. 5.**
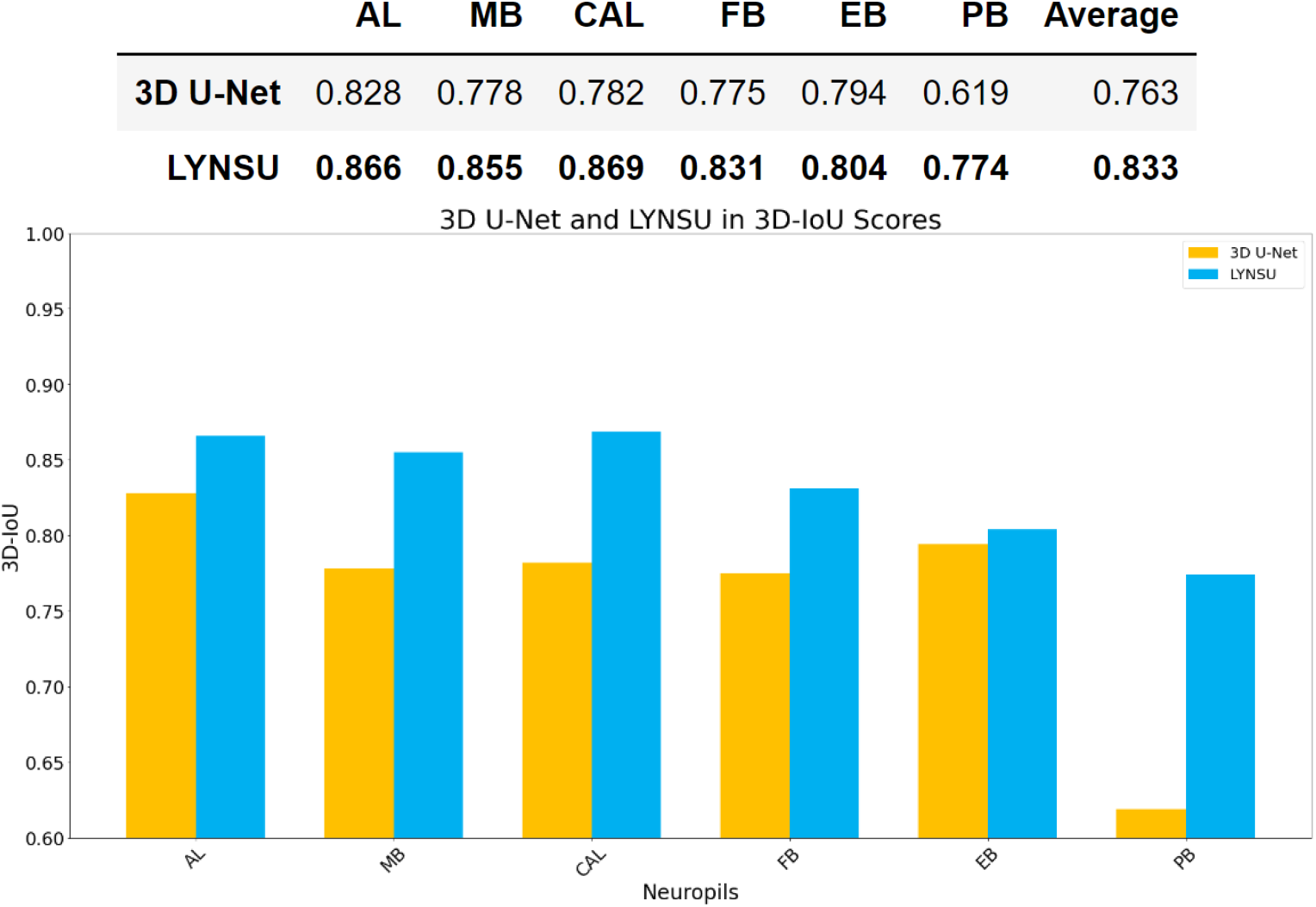
3D IOU scores for the segmentation of six neuropils or brain structures, compared between LYNSU and 3D U-net (without YOLO), on the test set. The former outperform the latter in all tests.

## Methods and Materials

### Dataset

The 3D fluorescence image data we used in this study were obtained from the *FlyCircuit* database (http://www.flycircuit.tw), which hosts images from 28,573 *Drosohpila* brains. These *Drosophila* brain images were acquired using high-resolution confocal microscopy. Each image contains a channel of GAL4 signals for individual neurons and a channel of anti-DLG staining for neuropils, and signal of each voxel is represented by a value ranging from 0 to 255. The calibrated voxel size of the images is x:y:z = 0.32 × 0.32 × 1 μm, and the XY plane resolution is 1024×1024.

Two datasets were created from the FlyCircuit database for the training purpose. For the first dataset, we first randomly selected 1000 3D images from the FlyCircuit database to train the YOLOv7 model. We used the Labelimg tool to annotate the neuropil bounding boxes, with 440 images going through this annotation process. Afterward, we randomly selected 400 of these images to serve as the training set, while the remaining 40 images were used as the test set.

The second dataset was primarily prepared for the training of the 3D U-Net model. We randomly extracted 18 Drosophila brain images from our database, each representing a different type of Gal4 Driver. This approach ensured that our study’s dataset encompassed a variety of brightness characteristics. Next, the six target neuropils or structures, AL, MB, CAL, FB, EB, and PB, in the 18 brains were manually segmented by trained annotators using the ZEISS arivis Cloud tool. On average, the annotation of each neuropil took about 4 hours, meaning that we invested approximately 432 hours in this annotation work. We specifically requested that multiple annotators label the same brain to ensure consistency among multiple annotators. We verified the consistency of the human annotation and found that the 3D IoU scores among different annotators for the same brain reached 0.85, effectively ensuring the quality and consistency of our annotation work.

### Neuropils Detection and Localization

We adopted a two-stage model training strategy to achieve superior neuropil segmentation effects and efficient computational speed. This strategy significantly enhances the accuracy of segmentation and effectively reduces the GPU computational resource consumption. We used the NVIDIA A100 40GB GPU in the first stage for model training. These images were projected into 2D images on the XY plane using two different projection methods: summation of brightness along the Z-axis and maximum brightness value along the Z-axis. We trained the YOLOv7 model on the training set (400 images from the first dataset) through 100 iterations. On the test set (40 images), this model performed excellently, achieving a mean Average Precision (mAP)@0.5 of 0.9955, and within the range of mAP@0.5:0.95 (different IoU thresholds from 0.5 to 0.95 in steps of 0.05), it scored 0.8489. Note that due to spatial overlap between neuropils or structures, some of them share the same ROI. Specifically, MB and CAL share the same ROI, while FB, EB, and PB share the central complex’s ROI, and the AL neuropil has its own ROI. Therefore, in the resent study, we only train YOLOv7 to detect three different ROIs.

Next, we used YOLOv7 to perform detection on the remaining 560 images that were not involved in the training. We visually inspect the result and label a detection as success if the resulting ROI (bounding box) covers the entire neuropil. YOLOv7 achieves a high success rate of 99.33% and it takes only about ten milliseconds to detect a neuropil in each image.

### Neuropils Segmentation

For neuropil segmentation, we have 18 manually segmented brains (the second dataset) as the training and test set. We first use YOLOv7 to detect the ROIs from these brains. Subsequently, we extracted the corresponding 3D images from the ROIs. For effective data augmentation, we implemented zero-padding along the Z-axis, extending it to 124 layers, while downscaling the XY plane by reducing the resolution of the XY plane to 168×168.

We adopted an overlapping sliding window strategy for data augmentation by setting the window size to 64×128x128, with a sliding distance of 20 each time. We performed a mirroring process on the Z-axis to further increase data diversity. Taking the MB as an example, we generate 144 cubes for each set of neuropils, meaning that 16 brains can produce 2304 cubes as the training data for the 3D U-Net. We independently trained a 3D U-Net model for each unique neuropil in the second stage. The learning rate of these models was set to 0.0001, and Adam was chosen as the optimization algorithm. Regarding loss function selection, we adopted a composite loss function, combining Dice loss and Categorical Focal Loss.

Dice loss is a function specifically designed for image segmentation problems and particularly suitable for scenarios with class imbalances. The mathematical formula for Dice loss is:

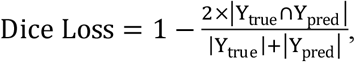

where Y_true_ is the set of ground truth labels and Y_pred_ is the set of labels predicted by the model. This formula can also be represented as:

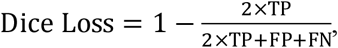

where TP represents true positive, FP false positive, and FN false negative. We set the weight ratio for the background and neuropils in the Dice loss as 0.4 and 0.6, respectively. We further incorporate weighting on category by setting the background weight *w*_*1*_ and neuropil weight *w*_*2*_. Thus, the weighted Dice loss function can be expressed as:

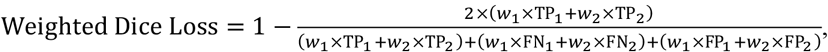

where TP_1_, FN_1_, FP_1_ are the true positive, false negative, and false positive for the background category, while TP_2_, FN_2_, FP_2_ are the corresponding quantities for the neuropil category. Through this weighting approach, we increased the relative importance of the neuropils (as opposed to the background) in the loss function, thereby making the model more focused on accurately segmenting neuropils.

Categorical Focal Loss (CategoricalFocalLoss) is primarily suitable for addressing class imbalance in multi-class classification problems. It’s defined by

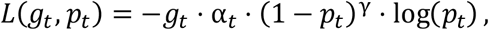

where *t* presents the class (neuropil or background), *g*_*t*_ *=* 1 if *t* is the ground truth class, *g*_*t*_ *=* 0 otherwise, *p*_*t*_ is the predicted probability for the class *t, α*_t_ is the balancing factor for *t*, and γ is a parameter that adjusts the predicted probability.

For the background class, which is account for the majority of the voxels, the model tends to predict a high probability *p*_*t*_. In this case, by increasing the γ parameter, we can reduce the loss contribution of these easily classified voxels. Conversely, for minority classes (neuropils), since *p*_*t*_ is generally lower, CategoricalFocalLoss does not excessively reduce the loss for these samples. This operation balances the loss between the background class and neuropils, reducing the impact of a higher number of predicted background class instances on the neuropil category, thereby enhancing the model’s ability to recognize imbalanced data.

In summary, the total loss function (Total Loss) is given by

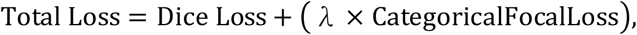

where λ is a hyperparameter used to balance the impact of the two loss functions, with a default value of 1, indicating equal consideration of both loss functions.

During the model training process, we set the batch size to 2 and the number of training epochs to 10. This ensures the model has sufficient data to learn from while avoiding overfitting. Additionally, to capture the model’s performance more accurately at different stages of training, whenever a lower loss value is observed on the validation set, we save the model’s current state. Such a strategy allows us to retain the best model during training and reduces the risk of model overfitting. By combining our chosen loss function, batch size, training epochs, and checkpoint monitoring strategy, we successfully designed a model training framework that converges quickly and has high generalization capabilities.

We conducted independent model training for each neuropil and create six LYNSU models for the six neuropils. Notably, each model only required approximately 20 minutes of training. In the inference phase, a model took an average of only 1.2 seconds to segment one neuropil.

## Discussions

This study successfully developed a novel, efficient, and accurate LYNSU segmentation workflow, specifically for neuropil segmentation in fluorescence images of *Drosophila* brains. By combining YOLOv7 and 3D U-Net, LYNSU substantially outperformed FCN and 2D U-Net, which were inadequate for the task. LYNSU also exhibited marked improvement over 3D U-Net by up to 15.5% in 3D IoU. It efficiently segmented a brain neuropil in just 7 seconds, demonstrating its suitability for large-scale databases.

This new method can potentially solve pressing issues in current connectomics, especially regarding spatial errors and computational efficiency in image alignment and registration. Previous methods often required aligning images using a standard brain template. Although being convenient, these methods unavoidably introduce spatial errors. Alternatively, one can segment neuropils or brain structures manually for individual images without using the standard brain template to achieve higher spatial accuracy. But manual segmentation is extremely time consuming and is not suitable for large-scale studies. In contrast, our algorithm combine the advantages of both approaches and achieves high spatial accuracy and temporal efficiency. Specifically, our algorithm can complete a neuropil segmentation task in just 7 seconds, which would take a human expert 4 hours. This breakthrough is significant considering the need for high-throughput connectomics research and will significantly accelerate the entire research process.

LYNSU offers several important applications. One can identify the innervated neuropils of a neuron more accurately because the neuropils can be identified in the same brain sample which the neuronal image is taken. Similarly, one can quantify the distribution of genes in different neuropils accurately. Finally, LYNSU can be used to study variability of neuropil morphology across individuals, which cannot be studied using the traditional methods that involving warping and registration images to the standard brain template.

## Statements and Declarations

The authors declare no conflict of interest.

## Acknowledgment

The authors would like to express their sincere gratitude to Dr. Hsiu-Ming Chang for his invaluable consultations regarding the database. Special thanks are also extended to the dedicated students who assisted in annotating brain images. This work was supported by National Science and Technology Council grant 111-2311-B-007-011-MY3, 112-2321-B-002-025 - and by the Brain Research Center under the Higher Education Sprout Project, co-funded by the Ministry of Education and the National Science and Technology Council in Taiwan.

